# Lateral Orbitofrontal Cortex and Basolateral Amygdala Regulate Sensitivity to Delayed Punishment during Decision-making

**DOI:** 10.1101/2022.04.13.488205

**Authors:** Anna E. Liley, Daniel B. K. Gabriel, Nicholas W. Simon

**Author notes:** Communicating Author: Nicholas W. Simon, Ph.D.

## Abstract

In real-world decision-making scenarios, negative consequences do not always occur immediately after a choice. This delay between action and outcome drives the underestimation, or “delay discounting”, of punishment. While the neural substrates underlying sensitivity to immediate punishment have been well-studied, there has been minimal investigation of delayed consequences. Here, we assessed the role of lateral orbitofrontal cortex (LOFC) and basolateral amygdala (BLA), two regions implicated in cost/benefit decision-making, in sensitivity to delayed vs immediate punishment. The delayed punishment decision-making task (DPDT) was used to measure delay discounting of punishment in rodents. During DPDT, rats choose between a small, single pellet reward and a large, three pellet reward accompanied by a mild foot shock. As the task progresses, the shock is preceded by a delay that systematically increases or decreases throughout the session. We observed that rats avoid choices associated with immediate punishment, then shift preference toward these options when the punishment is delayed. LOFC inactivation did not influence choice of rewards with immediate punishment, but decreased choice of delayed punishment. We also found that BLA inactivation reduced choice of delayed punishment, but this was only evident when punishment was initially delivered immediately after a choice, then preceded by a delay as the task progressed. Therefore, both LOFC and BLA contribute to the delay discounting of punishment, and may serve as promising therapeutic targets to improve sensitivity to delayed punishment during decision-making.

**Significance Statement:** Negative consequences occurring after a delay are often underestimated, which can lead to maladaptive decision-making. While sensitivity to immediate punishment during reward-seeking has been well-studied, the neural substrates underlying sensitivity to delayed punishment remain unclear. Here, we used the Delayed Punishment Decision-making Task to determine that lateral orbitofrontal cortex and basolateral amygdala both regulate the discounting of delayed punishment, suggesting that these regions may be potential targets to improve decision-making in psychopathology.

## Introduction

Many psychiatric diseases are characterized by insensitivity to detrimental outcomes (Bechara, 2005; Hartley and Phelps, 2012; Jean-Richard-dit-Bressel et al., 2019; Orsini and Simon, 2020). One factor that drives this insensitivity is the presence of a delay that often precedes occurrence of these outcomes (Murphy et al., 2001; Bechara and Dolan, 2002; Field et al., 2019). For example, individuals with substance misuse problems seek out drugs to receive immediate positive reinforcement, but often fail to consider impending withdrawal symptoms or financial/legal concerns because they occur later in time. This can be attributed to “delay discounting”, wherein the motivational value of delayed outcomes is underestimated compared to immediate outcomes.

While delay discounting of rewards has been well-studied (Floresco et al., 2008; Kable and Glimcher, 2010; Mar et al., 2011; Bickel et al., 2014; Burton et al., 2014; Frost and McNaughton, 2017), there is minimal research investigating the neurobiological mechanisms underlying sensitivity to immediate vs delayed punishment. Furthermore, preclinical research on punished reward-seeking has primarily focused on consequences that occur immediately after an action (Pollard and Howard, 1979; Jonkman et al., 2012; Orsini et al., 2015a; Park and Moghaddam, 2017; Jean-Richard-Dit-Bressel et al., 2018; Halladay et al., 2020; but see Rodríguez et al., 2018). To address this discrepancy, we developed the rat Delayed Punishment Decision-making Task (DPDT), which offers choice between a small reward and a large reward followed by a mild foot shock. The shock initially occurs immediately after choice, but is preceded by a systematically escalating delay as the task progresses (Liley et al., 2019). Rats initially avoid the punished option, then shift preference toward the punished option as delays increase, thereby demonstrating delay discounting of the negative motivational value as a function of delay. Moreover, despite comparable decision-making with immediate punishment, males discount delayed punishment to a greater degree than females. Critically, delay discounting of punishment is not correlated with discounting of rewards, suggesting that these processes may be governed by distinct neurobiological mechanisms (Liley et al., 2019).

The lateral orbitofrontal cortex (LOFC) and basolateral amygdala (BLA), two brain regions with dense reciprocal connections (Price, 2007), are two candidates that likely drive discounting of delayed punishment during reward-seeking. OFC is a prefrontal cortical brain region that receives input from all major sensory systems in addition to influences from limbic regions (Mcdonald, 1991; Carmichael and Price, 1995), making it an optimal site for integration of perceptual and emotional information to guide decision-making. LOFC is involved with a number of cognitive processes that are critical for cost-benefit decision-making (Padoa-Schioppa and Assad, 2006; Karimi et al., 2019), including reward/punishment integration (Sara E. Morrison, Alexandre Saez, Brian Lau, 2012; Orsini et al., 2015b; Jean-Richard-Dit-Bressel and McNally, 2016), and delayed reward discounting (Mobini et al., 2002a; Roesch et al., 2006; Zeeb et al., 2010); both of which are critical components of delayed punishment driven decision-making. Moreover, BLA regulates punishment-induced suppression of reward seeking and is involved with attribution of salience to aversive cues (Jean-Richard-dit-Bressel and McNally, 2015; Piantadosi et al., 2017; Hernandez et al., 2019). BLA is also involved with goal-directed behavior and encodes outcome-specific values to guide actions (Wassum and Izquierdo, 2015). Additionally, it associates cues with rewards during events, making this region imperative for utilizing past experiences to flexibly guide decisions for optimal outcomes (Schoenbaum et al., 2000). Thus, both regions likely contribute to integrating delays with punishment to guide decision-making.

Here, we separately assessed the involvement of LOFC and BLA in sensitivity to delayed punishment during decision-making using pharmacological inactivation of each region prior to DPDT testing. We also compared the effects of LOFC and BLA inactivation between male and female rats. Finally, to test if effects of inactivation were influenced by task design or impaired behavioral flexibility, LOFC and BLA were individually inactivated prior to a modified version of DPDT with descending punishment delays (REVDPDT).

## Methods

### Subjects

73 Long Evans rats obtained from Envigo were aged 70 days upon arrival and used for these experiments (Total LOFC: n = 38, Female: 18, Male: 20; Total BLA: n = 35, Female: 15, Male: 20). Rats were restricted to 85% free feeding weight one week prior to behavioral training to encourage motivation and pursuit of rewards during task performance. Free-feeding weights were altered throughout the experiment in accordance with Envigo growth charts to account for growth. All rats were individually housed and maintained on a 12-hour reverse light/dark cycle. All methods were approved by the University of Memphis Institutional Animal Care and Use Committee.

### Surgery

Prior to behavioral assessments, rats underwent bilateral cannulation surgery for either LOFC (3.0 mm AP, 3.2 ML, and 4.0 DV from skull surface (Roesch et al., 2006; Bjerke et al., 2019)) or BLA (2.8 mm AP, 5.0 ML, and 8.7 DV from skull (Orsini et al., 2015b; Bjerke et al., 2019)) infusions. Rats were anesthetized in an isoflurane gas induction chamber, then placed into a stereotaxic apparatus (Kopf; Tujunga, CA) while resting on a heating pad adjusted to 40 degrees C. Isoflurane was provided throughout surgery via a nose cone. Cannulae were held in place by a dental cement headcap anchored by three bone screws. Upon completion of surgery, rats were subcutaneously given 1mL of sterile saline, and a solution of Acetaminophen and H2O along with a dish of moistened food during recovery. Rats were closely monitored for signs of infection or distress during the next week, with cage bedding changed daily for the first 3 days.

### Behavior Apparatus

Testing was conducted in standard rat behavioral test chambers (Med Associates; Fairfax, VT) housed within sound attenuating cubicles. Each chamber was equipped with a recessed food pellet delivery trough fitted with a photo beam to detect head entries, and a 1.12-watt lamp to illuminate the food trough. Food pellets were delivered into the food trough 2 cm above the floor centered in the side wall. Two retractable levers were located on the left and right side of the food trough 11 cm above the floor. A 1.12-watt house light was mounted on the opposing side wall of the chamber. Beneath the house light was a circular nose poke port equipped with a light and photo beam to detect entry. The floor of the test chamber was composed of steel rods connected to a shock generator that delivered scrambled foot shocks. Locomotor activity was assessed throughout each session with infrared activity monitors located on either side of the chamber just above the floor. Test chambers are interfaced with a computer running MedPC software, which controlled all external cues and behavioral events.

### Shaping Procedures

Food restriction and behavioral training began after one week of recovery. Prior to acquisition of DPDT, rats underwent a series of shaping procedures. Rats were first taught to associate the food trough with food pellets during magazine training. In separate sessions, rats then trained to press a single lever (left or right, counterbalanced across groups) to receive one pellet of food. After performing 50 reinforced lever presses within 30 minutes, rats trained to press the opposite lever under the same criterion. Following this were shaping trials in which both left and right levers were retracted, and rats were required to nose poke into the food trough during a period of illumination from both the house and food trough lights. Nose poking evoked the extension of a single lever (either left or right in pseudorandom order). Each subsequent lever press was reinforced with a single pellet, along with extinguishing of house and trough lights and retraction of the lever. After achieving a minimum of 30 presses of each lever in a 60-minute time span, rats progressed to magnitude discrimination training.

The 30-minute reward magnitude sessions utilized 2 levers with counterbalanced presses producing either 1 or 3 pellets. As in the previous stage of training, each trial began with the illuminated house and trough lights, after which a nose poke into the trough led to extension of one or both levers. A press on one lever produced a single pellet while the other produced 3, followed by lever retraction, termination of all cues, and progression to a 10 ± 2s intertrial interval (ITI). There were 5 blocks of 18 trials in each task, with the first 8 providing only a single lever (forced choice) the next 10 both levers (free choice). Once rats achieved >75% preference for the large reward during free choice trials, they began either DPDT or REVDPDT training.

### Delayed Punishment Decision-making Task (DPDT)

During DPDT, rats chose between a small reward and larger reward associated with punishment preceded by varying delays. DPDT methodology was comparable to magnitude discrimination described above, with choice between small and large food pellet reinforcers. However, in this task the large option was accompanied by a mild, one second foot shock. This shock either initially occurred immediately after a choice, then systematically took place later in time throughout the task or vice versa (Figure 1).

**Figure 1.**
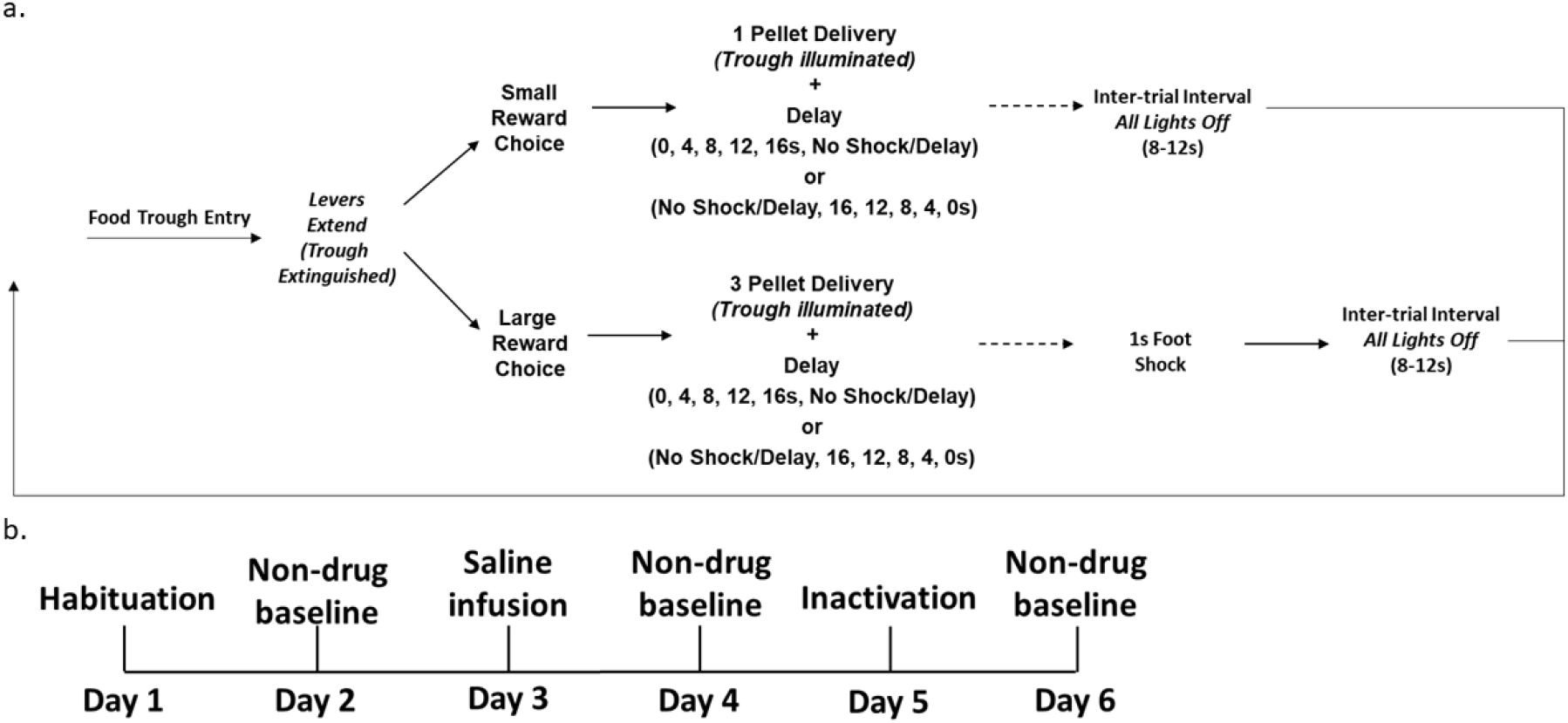
a) Delayed Punishment Decision-making Task (DPDT and REVDPDT). Rats chose between two levers, one delivering a one-pellet reward and the other delivering a three-pellet reward accompanied by a delayed foot shock (delay sequence: 0, 4, 8, 12, 16s, No Shock/Delay for DPDT; No Shock/ Delay, 16, 12, 8, 4, 0s for REVDPDT). b) Six-day micro-infusion schedule for LOFC and BLA. Inactivation drug cocktail and sterile saline were counterbalanced across subjects.

Trials began with illumination of the house light and food trough, after which a nose poke into the trough caused one or both levers to extend simultaneously. A press on one lever dispensed a single pellet, while the other dispensed three pellets with a one-second mild foot shock. After all outcomes were delivered, the house light extinguished, and the next trial proceeded after an ITI of 10 ± 2s. The sessions were divided into 6 blocks, with 2 forced choice and 10 free choice trials in each block for a total of 72 trials. The first two trials of each block were “forced choice” trials in which only a single lever was available to establish the reward/punishment parameters within that block. The following 10 trials were “free-choice” trials in which both levers extended, allowing rats to choose a preferred lever. During the first block, shock occurred immediately after lever press. In each subsequent block, a delay was introduced preceding shock that extended from 0, 4, 8, 12, and 16 seconds, followed by a block with and no shock/delay (Figure 1). Notably, on trials in which the unpunished lever was chosen, the ITI increased by a period equivalent to the delay preceding shock in that block (4, 8, 12, 16s) to maintain consistency of trial length regardless of choice.

To minimize individual differences in performance and allow sufficient parametric space for choice to increase or decrease following manipulations, shock intensity was titrated for each individual rat until: 1) preference for the punished reward was between floor (0% choice of punishment) and ceiling (100% choice of punishment), and 2) there was either a positive slope between blocks 1 and 6 of DPDT or a negative slope between blocks 1 and 6 of REVDPDT, indicative of discounting of delayed punishment. Shock intensity began at 0.05 mA, then increased by .02, .03, .05, or 0.1 mA (based on sensitivity) in subsequent sessions if rats completed >85% of trials. This incremental increase in shock intensity limited omissions and allowed rats to acquire task parameters. Upon reaching the final shock intensity, subjects trained until they achieved stability, which consisted of no more than a 10% overall shift in daily choice behavior for 2-3 days.

### DPDT with descending delays (REVDPDT)

To confirm that effects of inactivation were not driven by task design or aberrant behavioral flexibility, a subset of subjects were trained in a reversed task (REVDPDT). This task was identical to DPDT, except the delays were presented in descending order, beginning with no shock/delay, 16, 12, 8, 4, then 0 seconds preceding foot shock (Figure 1). Criteria for stability and shaping procedures were comparable to those used for DPDT.

### LOFC and BLA Inactivation

After rats reached stable performance in DPDT or REVDPDT, they underwent habituation sessions to acclimate to infusion procedure handling. They then received bilateral drug micro-infusions to inactivate either bilateral LOFC or BLA. A drug cocktail of GABA agonists baclofen (Reis and Duarte, 2006) and muscimol (Chandra et al., 2010) dissolved in sterile saline (concentration: 250ng/μl, .5 μl infusion volume over 1 min (Piantadosi et al., 2017b; Orsini et al., 2018)) was administered into each hemisphere via an automated infusion pump and 2 50μl Hamilton syringes. Behavioral testing commenced after a 15-minute absorption period. After a day of baseline testing with no treatment, subjects were given bilateral sterile saline micro-infusions (5μl infused at .5μl/min). Drug/saline order was counterbalanced across subjects.

### Histology

Rats were euthanized with Euthasol, and perfusions were conducted with saline and 10% formalin solution. Brains were extracted, stored in 10% formalin solution, sliced at 60-100 μm using a Cryostat, and mounted onto slides. Cannula placements and infusion localization were confirmed via light microscopy (Bjerke et al., 2019)

### Experimental Design and Statistical Analysis

Custom-made MATLAB scripts were used to compile behavioral data, and all statistical analyses were conducted using IBM SPSS Statistics 24. If Mauchly’s test of sphericity was violated, Greenhouse-Geisser values and degrees of freedom were used accordingly. If a rat failed to make any choices during a block of the task, the slope of that subject’s curve was used to extrapolate that missing data point. If two or more blocks of behavioral data were missing, that rat was removed from analysis due to excessive omissions.

Following task acquisition, stable decision-making for DPDT and REVDPDT were measured using a day x block repeated measures ANOVA, quantified as lack of effect of day and a significant effect of block. Effects of micro-infusions on behavior were analyzed via sex x infusion (drug vs saline) x block ANOVA. Latency to lever press during testing was evaluated using a mixed sex x safe vs punished lever ANOVA.

## Results

### DPDT/REVDPDT Acquisition and Stability

The mean number of days to complete shaping procedures prior to DPDT (magazine training, FR1 for both levers, nose poke, and magnitude discrimination training) was 8.56 for females and 6.89 for males, with no significant difference between sexes (*t* (26) = 1.368, p = .183). Females required significantly more training sessions to reach stability on DPDT (female mean = 38.67 days, male mean = 14.53 days; *t* (26) = 3.15, p = .004). After successful training, there were significant effects of block for all subjects in DPDT, such that subjects shifted choice preference from the safe reward to the punished reward as punishment delay increased (LOFC group: *F*_(2.159, 21.587)_ = 29.304, *p* < .001; BLA group: *F*_(2.583, 28.409)_ = 13.524, *p* < .001). There were no significant differences between sexes in DPDT performance (LOFC: *F*_(5, 50)_ = .627, *p* = .680; BLA group: *F*_(5, 55)_ = 2.204, *p* = .067).

A separate group of rats trained in REVDPDT, in which punishment delays were presented in descending instead of ascending order (Figure 1). There was no difference in length of shaping for REVDPDT between males and females (female mean = 9.27 days, male mean =7.61 days; *t* (27) = 1.618, *p* = .117). Unlike in DPDT, there were no sex difference in sessions required to achieve stability for REVDPDT (female mean= 30.36 days, male mean = 30.67 days; *t* (27) = .03, *p*=.976). After task acquisition, there were significant effects of block (LOFC group: *F*_(2.716,27.164)_ = 18.059, *p* < .001; BLA group: *F*_(5, 45)_ = 11.075, *p* < .001), such that rats shifted choice away from the punished option as punishment became less delayed/more proximal to the action. There were no sex differences in REVDPDT (LOFC group: *F*_(5, 50)_ = .657, *p* = .657; BLA group = (*F*_(5, 45)_ = 1.191, *p* = .329).

### Effects of LOFC Inactivation on DPDT

We assessed the effects of acute pharmacological LOFC inactivation on sensitivity to delayed punishment prior to DPDT using 9 male and 3 female rats, with bilateral cannulae placement in LOFC confirmed before any analyses (Figure 2a). There was a main effect of block (*F*_(2.793, 27.931)_ = 26.736, *p* < .001; Figure 3a), such that rats chose the punished reward more frequently when punishment was delayed. There was no effect of sex (*F*_(1,10)_ = .018, *p* = .897), inactivation x sex interaction (*F*_(1, 10)_ = 1.024, *p* = .335), or inactivation x block x sex interaction (*F*_(5, 50)_ = .663, *p* = .653), so males and females were merged for all analyses (Figures 3b-c).

**Figure 2.**
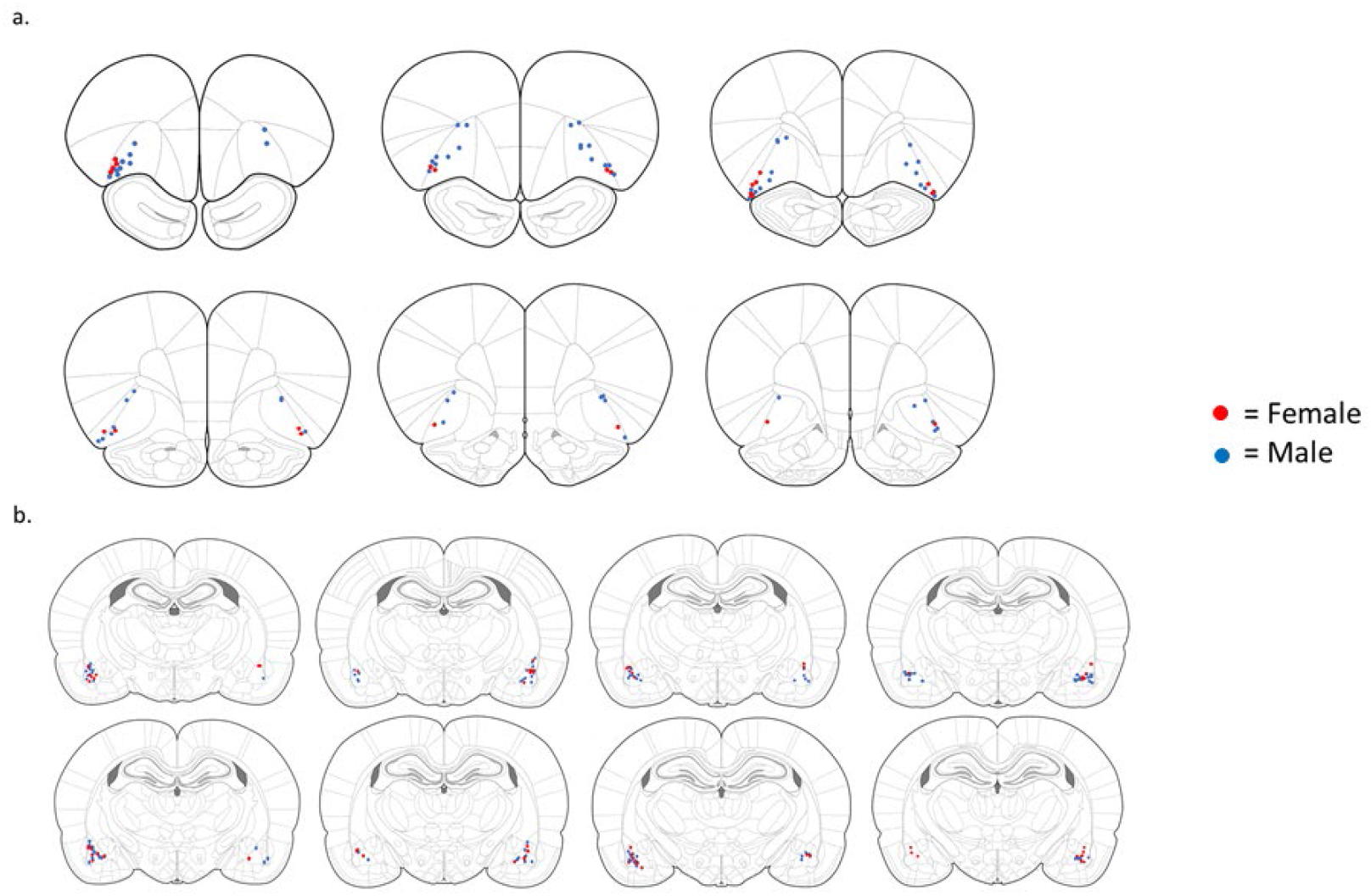
Histological confirmation of cannulae placements in LOFC and BLA.

**Figure 3.**
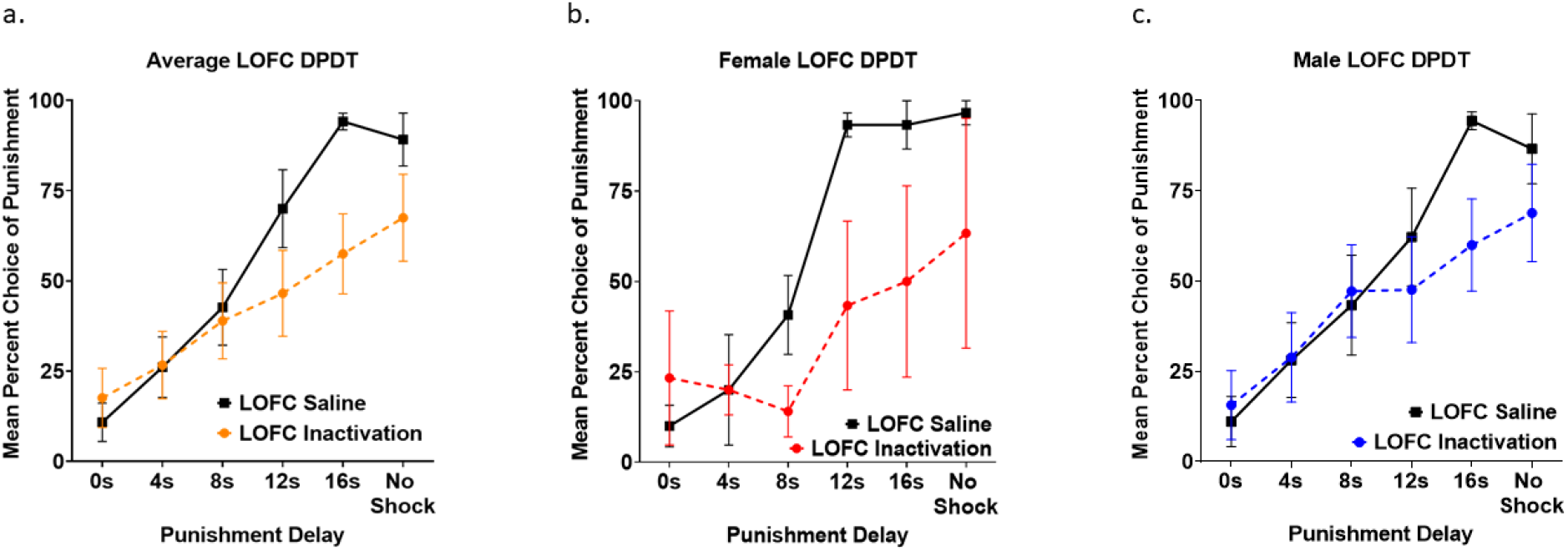
a) LOFC inactivation reduced choice of rewards with delayed punishment without affecting choice of immediate or short-delay punishment. b-c) Females and males showed comparable reduction in choice of delayed but not immediate punishment after LOFC inactivation. All panels display data as mean ± SEM.

OFC inactivation reduced overall choice of the punished reward (*F*_(1, 10)_ = 5.888, *p* = .036). Critically, there was also an inactivation x block interaction (*F*_(5, 50)_ = 3.261, *p* = .013; Figure 3a), such that LOFC inactivation only reduced large reward choice when punishment occurred after long delays, but not when punishment occurred immediately or after a short delay. Further investigation using two-tailed paired samples *t*-tests (see Table 1a) revealed no effects of inactivation in the first three blocks, a near significant difference between drug vs saline for the 12 second delayed shock (*p* = .074), and a significant difference for the 16 second delayed shock (*p* = .005). Finally, there was no effect of LOFC inactivation during the final, unpunished block (*p* = .16), suggesting that LOFC inactivation did not cause gross motivational deficits or inability to discriminate reward magnitude.

**Table 1.** a) t-test Results Comparing Average Selection of the Punished Lever Following Inactivation vs Saline during LOFC DPDT. b) t-test Results Comparing Average Selection of the Punished Lever Following Inactivation vs Saline during LOFC REVDPDT. Both include mean differences between LOFC inactivation and saline.

| LOFC |  |  |  |  |  |
| --- | --- | --- | --- | --- | --- |
| a. | Delay | Mean Differences | t | df | p |
|  | 0s | 6.76 | 0.894 | 11 | 0.39 |
|  | 4s | 0.56 | 0.083 | 11 | 0.935 |
|  | 8s | -3.75 | -0.516 | 11 | 0.616 |
|  | 12s | -23.43 | -1.971 | 11 | 0.074 |
|  | 16s | -36.67 | -3.552 | 11 | 0.005* |
|  | No shock | -21.67 | -1.516 | 11 | 0.158 |
| b. | Delay | Mean Differences | t | df | p |
|  | No shock | -11.79 | -1.863 | 13 | 0.085 |
|  | 16s | -25.56 | -2.816 | 13 | 0.015* |
|  | 12s | -16.39 | -1.34 | 13 | 0.203 |
|  | 8s | -9.75 | -0.821 | 13 | 0.426 |
|  | 4s | -7.08 | -0.582 | 13 | 0.571 |
|  | 0s | -3.36 | -0.392 | 13 | 0.701 |

Next, we assessed the effects of LOFC inactivation on omitted trials during DPDT. There was a main effect of inactivation (*F*_(1, 10)_ = 10.494, *p* = .009; Figure 4a) such that omissions were greater following inactivation compared to saline infusions. There was also an effect of block (*F*_(1.706, 17.061)_ = 5.734, *p* = .015), with subjects omitting more trials early in the session wherein punishment had shorter delay times. There was no inactivation x block interaction (*F*_(1.590, 15.901)_ = 2.220, *p* = .148). There was a trend toward a main effect of sex in which females displayed more omissions throughout the task than males (*F*_(1, 10)_ = 4.395, *p* = .062; Figure 4b-c). There was also an inactivation x sex interaction (*F*_(1, 10)_ = 9.655, *p* = .011), with females showing increased omitted trials after LOFC inactivation compared to males.

**Figure 4.**
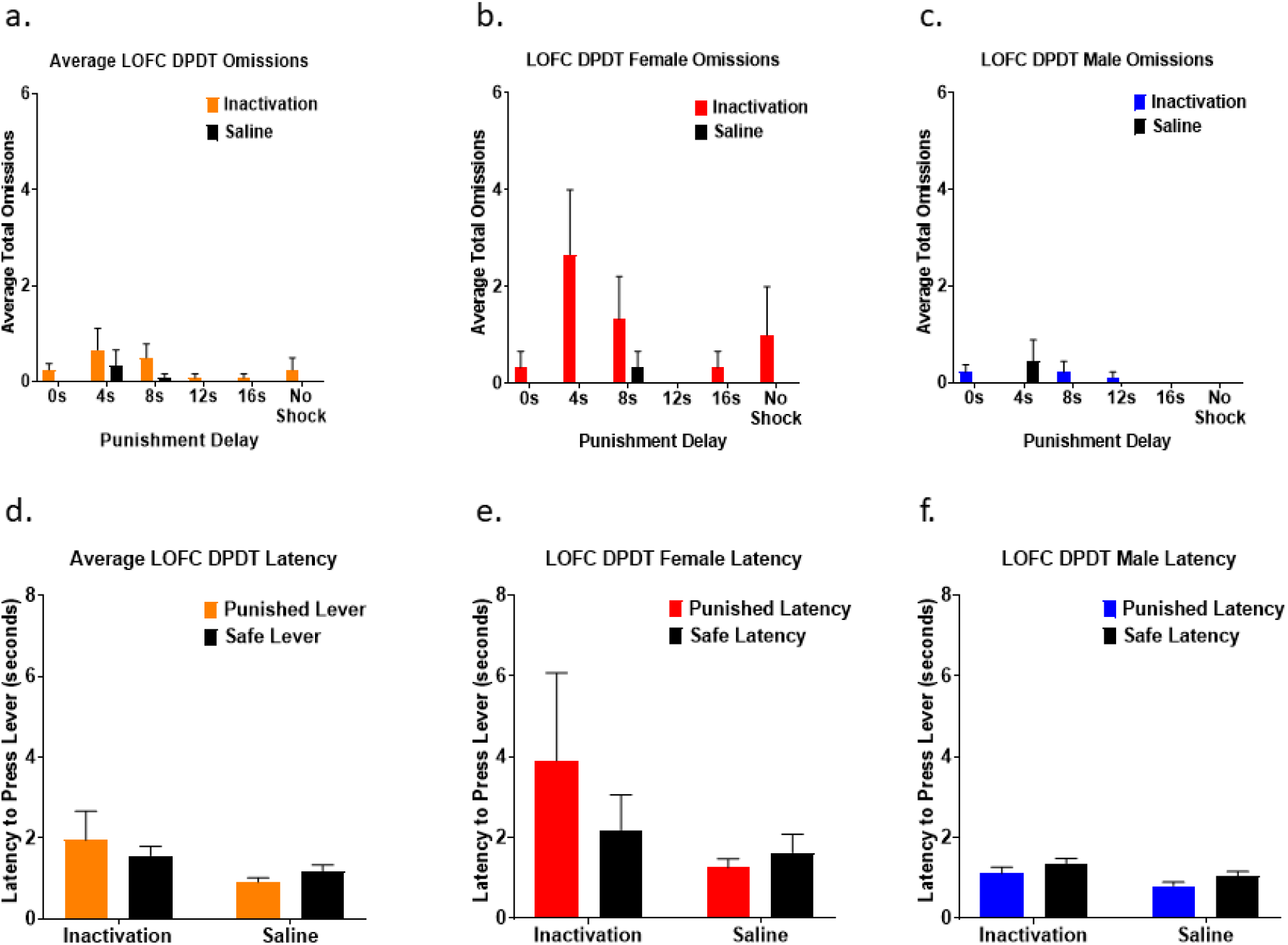
a) LOFC inactivation increased omissions compared to saline, and omissions were most prevalent when punishment occurred after shorter delays. b-c) Females displayed more omissions throughout the task than males. d) LOFC infusions did not affect latency to choose either lever. e-f) Females took longer to choose the punished lever after LOFC inactivation than males. All panels display data as mean + SEM.

We next investigated the effects of LOFC inactivation on latency to choose a lever. There was no significant difference in latency to choose safe vs punished levers (*F*_(1, 11)_ = .073, *p* = .792; Figure 4d), nor was there an effect of LOFC inactivation (*F*_(1, 11)_ = .003, *p* = .960). However, there was a trend toward an inactivation x lever type interaction (*F*_(1, 11)_ = 4.435, *p* = .059), such that inactivation lengthened the time required for subjects to choose the punished but not safe lever. There was also an effect of sex (*F*_(1, 11)_ = 8.871, *p* = .013; Figures 4e-f), with females taking longer than males to make a choice, but no sex x lever type interaction (*F*_(1, 11)_ = 2.319, *p* = .156).

### Effects of LOFC Inactivation on REVDPDT

To test that the effects observed with DPDT were not solely due to inflexible behavior (leading to a “flattened” discounting curve), we trained rats in a reversed version of DPDT (REVDPDT) with descending delays preceding punishment (blocks: no shock, 16s, 12s, 8s, 4s, 0s). Analysis of brain sections determined that 5 female and 9 male rats (n = 14) had accurate cannula placement in LOFC (Figure 2a). Importantly, as with DPDT, there was a significant main effect of block (*F*_(5, 60)_ = 21.468, *p* < .001; 5a), such that rats shifted choice away from the punished reward as delay decreased. There was no effect of sex (*F*_(1, 12)_ = 3.119, *p* = .103), inactivation x sex interaction (*F*_(1, 12)_ = .002, *p* = .961), or inactivation x block x sex interaction (*F*_(5, 60)_ = .381, *p* = .860), so males and females were again merged for subsequent analyses (Figure 5b-c).

**Figure 5.**
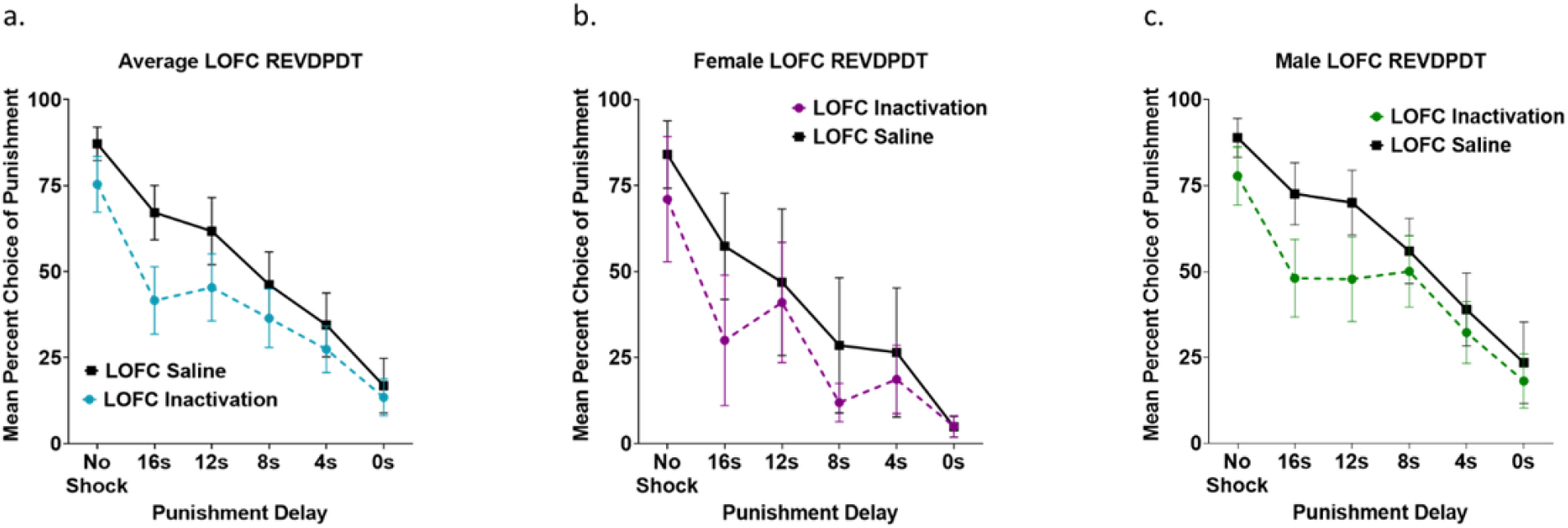
a) Inactivation of LOFC during REVDPDT shifted choice away from the punished reward as delays decreased. b-c) Females and males displayed similar reduction in choice of the punished lever compared to safe when delays decreased during LOFC inactivation. All panels display data as mean + SEM.

There were no effects of inactivation on choice of the punished option (*F*_(1, 12)_ = 1.920, *p* = .191), nor an inactivation x block interaction (*F*_(5, 60)_ = 1.128, *p* = .355). However, based on the LOFC inactivation exerting the most substantial effects during the 16 second delayed punishment in the standard DPDT (Figure 5a), we probed for an effect of inactivation on this block during REVDPDT. We observed that LOFC inactivation did indeed reduce choice of the punished reward in this block (*t* (13) = −2.816, p = .015), with no differences observed in other blocks (Table 1b). This revealed that, as in the original task, LOFC inactivation reduced choice of the punished reward when punishment occurred after a long (16s) delay, but not when punishment coincided with choice after shorter (0-12s) delays.

We next investigated the effects of LOFC inactivation on omissions during REVDPDT. This revealed a near significant effect of inactivation (*F*_(1, 12)_ = 4.690, *p* = .051), such that LOFC inactivation increased omissions compared to saline. There was also a significant effect of block (*F*_(2.363, 28.355)_ = 4.277, *p* = .019) with omissions increasing after the first, unpunished block (Figure 6a). There was no inactivation x block interaction (*F*_(2.760, 33.122)_ = 2.160, *p* = .116). There was an effect of sex (*F*_(1, 12)_ = 11.974, *p* = .005), with females omitting more trials than males. There was also an inactivation x sex interaction (*F*_(1, 12)_ = 4.887, *p* = .047), with females, but not males, demonstrating an increase in omissions after LOFC inactivation (Figure 6b-c).

**Figure 6.**
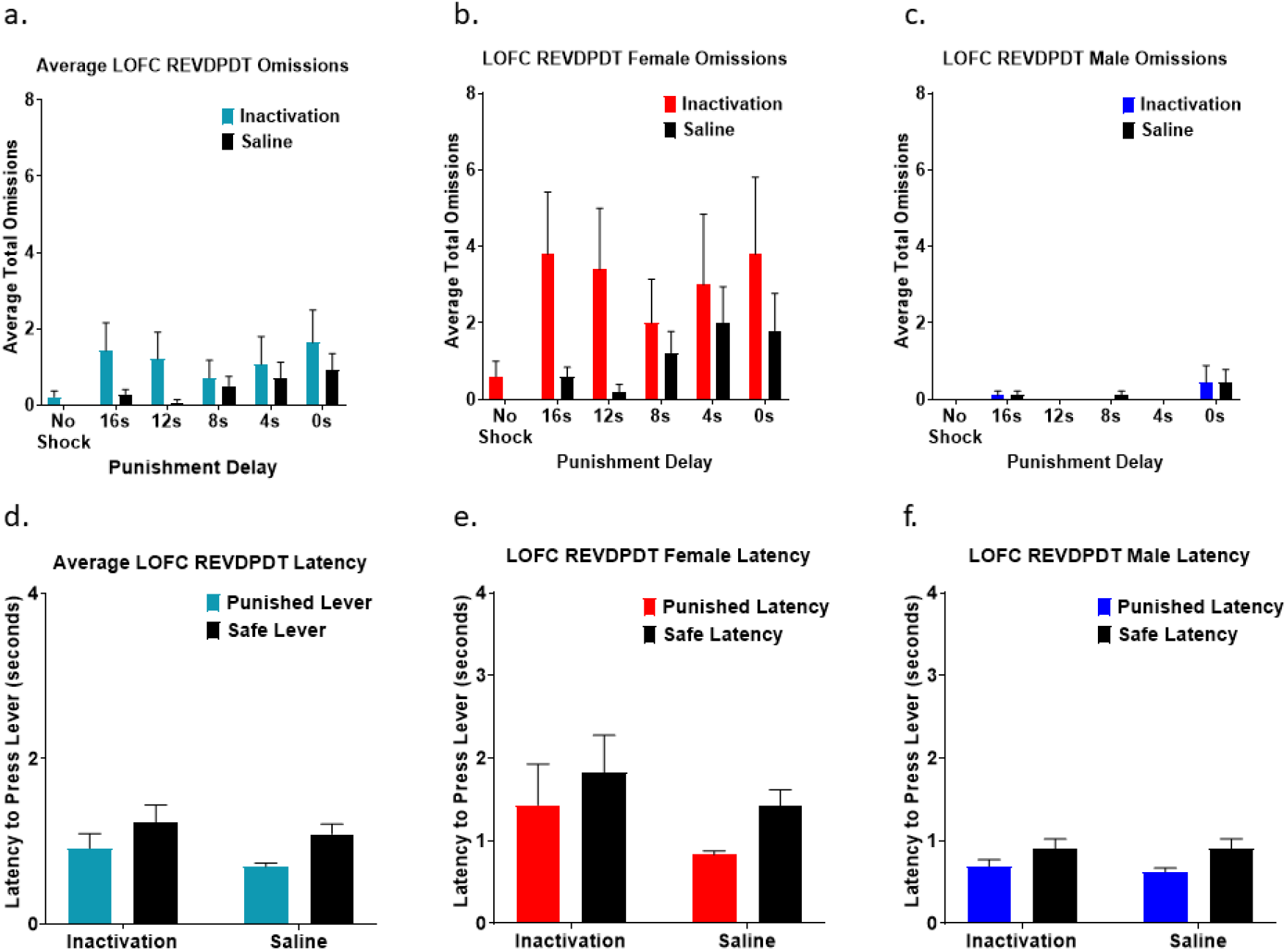
a) Inactivation of LOFC during REVDPDT increased omissions during inactivation compared to saline. b-c) Females omitted more trials than males throughout the task. d) There were no differences in latency to choose either the punished or safe levers for both inactivation and saline infusions during REVDPDT. e-f) Females required more time to select the punished lever than males. All panels display data as mean + SEM.

Finally, we examined the effects of LOFC inactivation on response latency. There was no effect of inactivation (*F*_(1, 11)_ = 3.403, *p* = .092) or inactivation x lever type interaction (*F*_(1, 11)_ = .300, *p* = .595; Figure 6d). There was a difference in time taken to choose a lever (*F*_(1, 11)_ = 11.619, *p* = .006). There was also a main effect of sex (*F*_(1, 11)_ = 11.420, *p* = .006; Figures 6e-f) such that females were slower to respond than males. However, there were no sex x lever type (*F*_(1,11)_ = 1.911, *p* = .194) or sex x drug interactions (*F*_(1,11)_ = 2.745, *p* = .126).

### Effects of BLA Inactivation on DPDT

Analysis of brain sections determined that 5 female and 9 male rats (n = 14) had accurate cannula placement (Figure 2b). As previously, there was a significant effect of block (*F*_(5, 60)_ = 16.312, *p* < .001; Figure 7a) indicating that rats selected the punished reward more frequently when punishment was delayed. There was no main effect of sex (*F*_(1, 12)_ = .231, *p* = .639), inactivation x sex interaction (*F*_(1, 12)_ = .420, *p* = .529), block x sex (*F*_(5, 60)_ = .1.479, *p* = .210), or inactivation x block x sex interaction (*F*_(5, 60)_ = 1.093, *p* = .374), so sexes were combined for all analyses (Figure 7b-c).

**Figure 7.**
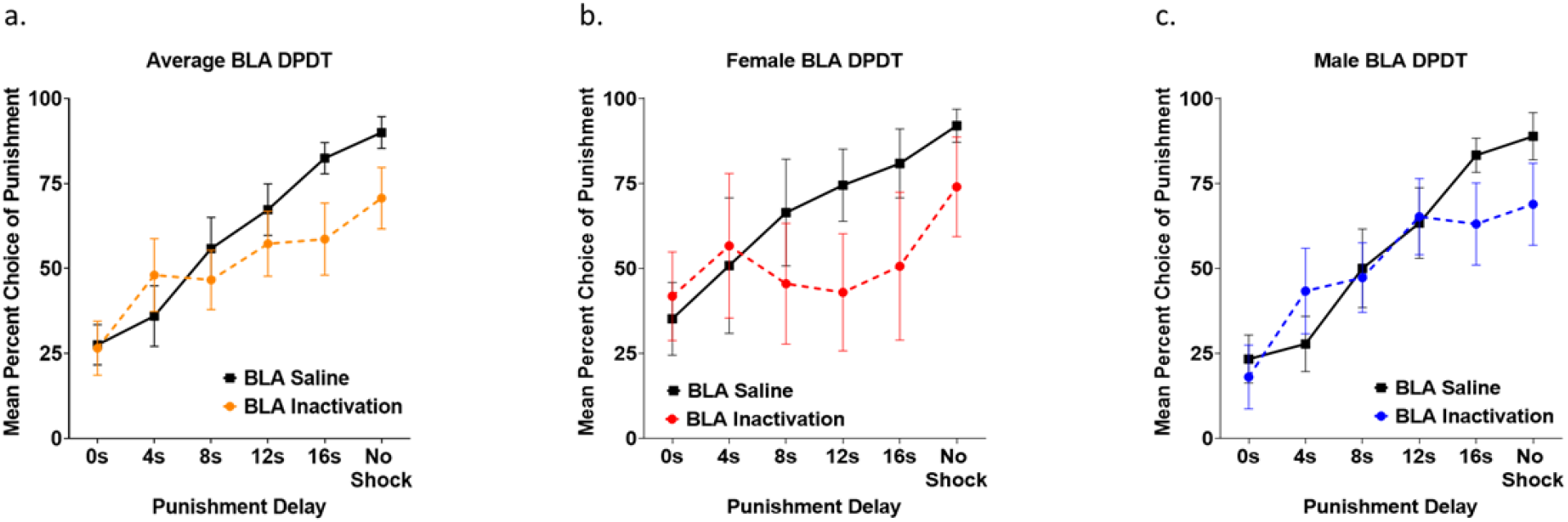
a) BLA inactivation did not affect choice when punishment was immediate, but reduced choice of the punished reward when delays were longer. b-c) Females and males both displayed reduction in choice of delayed punishment but not immediate punishment following BLA inactivation. All panels display data as mean + SEM.

There was no overall main effect of inactivation on choice (*F*_(1, 12)_ = 1.800, *p* = .205). However, there was an inactivation x block interaction (*F*_(5, 60)_ = 3.102, *p* = .015) such that BLA inactivation reduced choice of the punished reward when punishment was delayed (Figure 7a) but did not affect choice when punishment was immediate. Paired samples *t*-tests revealed this significant reduction in choice of the punished reward only occurred in the 16 second delay condition (*t* (13) = −2.787, p = .015; Table 2a). Interestingly, BLA inactivation also reduced choice of the large reward in the final, unpunished block (*t* (13) = −3.006, p = .010).

**Table 2.**
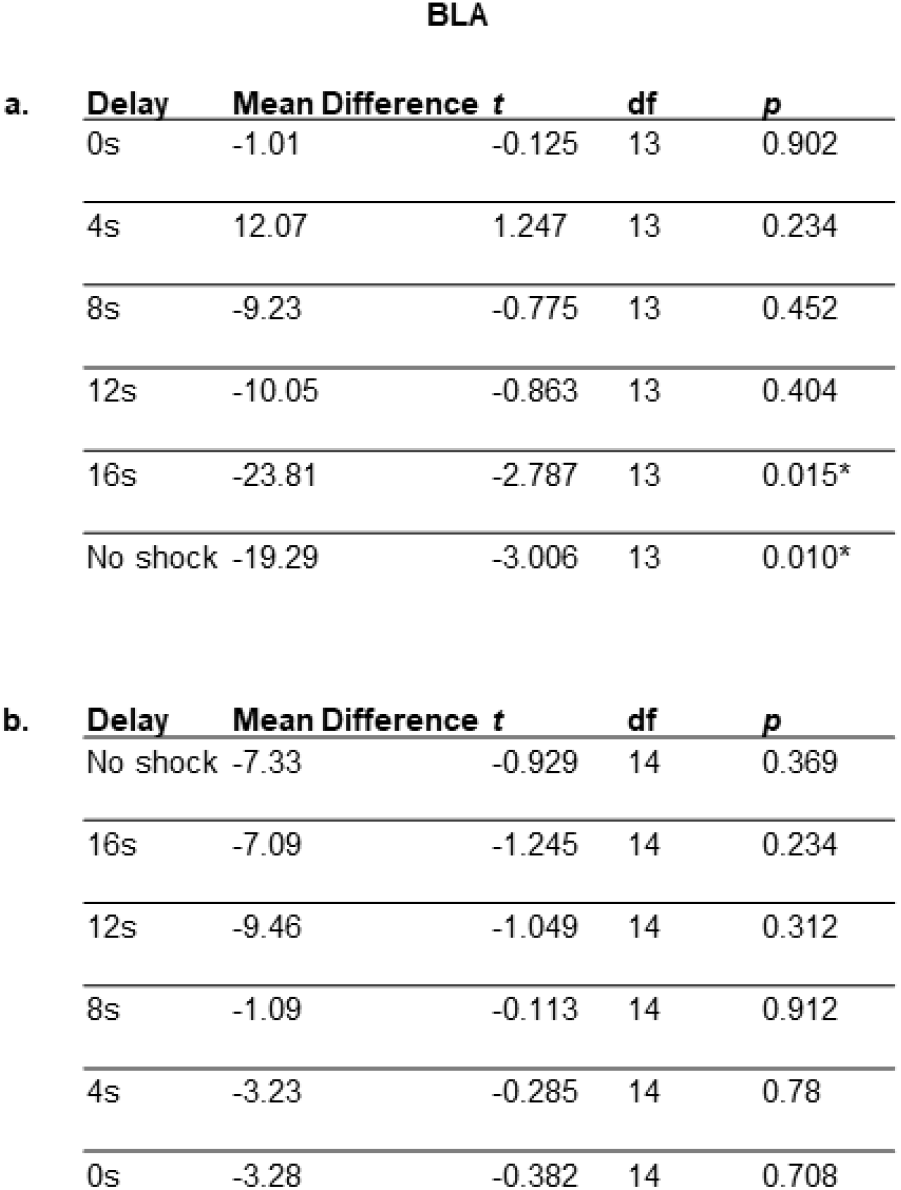
a) t-test Results Comparing Average Selection of the Punished Lever Following Inactivation vs Saline during BLA DPDT. b) t-test Results Comparing Average Selection of the Punished Lever Following Inactivation vs Saline during BLA REVDPDT. Both include mean differences between BLA inactivation and saline.

**BLA**
| **a.** | **Delay** | **Mean Difference** | **t** | **df** | **p** |
| --- | --- | --- | --- | --- | --- |
|  | 0s | -1.01 | -0.125 | 13 | 0.902 |
|  | 4s | 12.07 | 1.247 | 13 | 0.234 |
|  | 8s | -9.23 | -0.775 | 13 | 0.452 |
|  | 12s | -10.05 | -0.863 | 13 | 0.404 |
|  | 16s | -23.81 | -2.787 | 13 | 0.015* |
|  | No shock | -19.29 | -3.006 | 13 | 0.010* |

**Table 2.** a) t-test Results Comparing Average Selection of the Punished Lever Following Inactivation vs Saline during BLA DPDT.
| **b.** | **Delay** | **Mean Difference** | **t** | **df** | **p** |
| --- | --- | --- | --- | --- | --- |
|  | No shock | -7.33 | -0.929 | 14 | 0.369 |
|  | 16s | -7.09 | -1.245 | 14 | 0.234 |
|  | 12s | -9.46 | -1.049 | 14 | 0.312 |
|  | 8s | -1.09 | -0.113 | 14 | 0.912 |
|  | 4s | -3.23 | -0.285 | 14 | 0.78 |
|  | 0s | -3.28 | -0.382 | 14 | 0.708 |

There was an effect of block on omissions, such that subjects increased omissions as the session progressed (*F*_(1.260, 15.118)_ = 8.647, *p* = .007; Figure 8a). While there was no overall effect of BLA inactivation on omissions (*F*_(1, 12)_ = 3.470, *p* = .087), there was a near-significant inactivation x block interaction (*F*_(2.356, 28.276)_ = 3.095, *p* = .053), with BLA inactivation causing a greater increase in omissions as the task continued (Figure 8a). Additionally, there was an effect of sex in that females had more omissions than males (*F*_(1, 12)_ = 9.555, *p* = .009), but no inactivation x sex interaction (*F*_(1, 12)_ = .502, *p* = .492) or inactivation x block x sex interaction (*F*_(5, 60)_ = 1.891, *p* = .109; Figure 8b-c).

**Figure 8.**
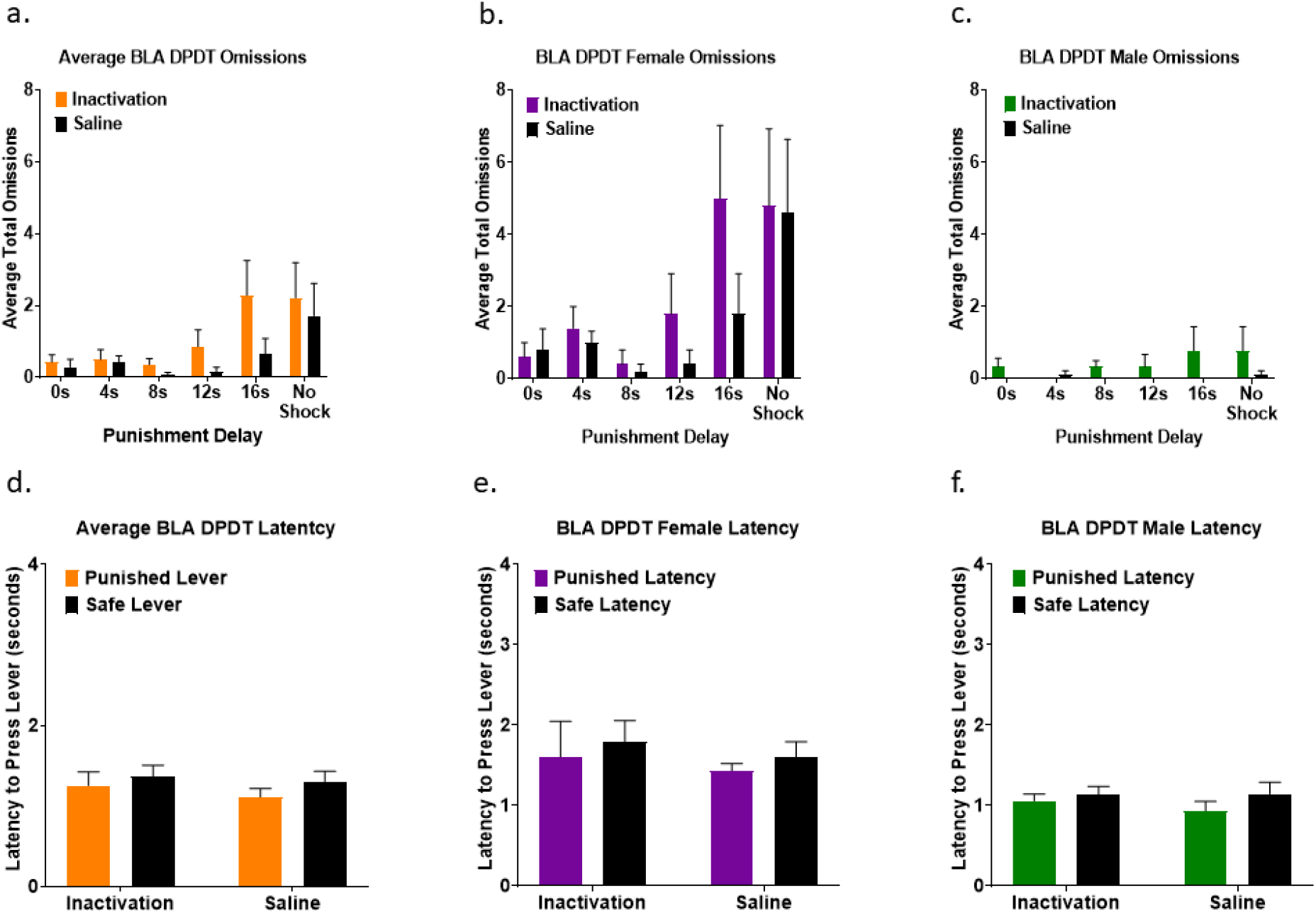
a) Subjects increased omissions during longer delay times of DPDT following BLA inactivation compared to saline. b-c) Females had more omissions than males throughout the task. d) There were no effects of inactivation or saline on latency to choose between the punished and safe levers. e-f) Females required more time to choose a lever than males. All panels display data as mean + SEM.

We next investigated the effects of BLA inactivation on decision latency. There was no main effect of inactivation (*F*_(1, 12)_ = .770, *p* = .397), latency between punished vs safe levers (*F*_(1,12)_ = 1.339, *p* = .270), or inactivation x lever type interaction (*F*_(1, 12)_ = .178, *p* = .680; Figure 8d). There was a main effect of sex (*F*_(1, 12)_ = 12.551, *p* = .004), with females taking longer to make a choice than males; but no sex x lever type interaction (*F*_(1,12)_ = .016, *p* = .903) or sex x inactivation interaction (*F*_(1,12)_ = .223, *p* = .645; Figure 8e-f).

### Effects of BLA Inactivation on REVDPDT

BLA was inactivated prior to REVDPDT in 6 female and 9 male rats (Figure 2b). As expected, there was a main effect of block (*F*_(5, 65)_ = 12.065, *p* < .001; Figure 9a) such that the initial preference for the large reward shifted toward the safe reward as punishment delay decreased. Similar to DPDT, there was no main effect of sex (*F*_(1, 13)_ = 2.489, *p* = .139), sex x inactivation interaction (*F*_(1, 13)_ = .015, *p* = .905), block x sex interaction (*F*_(5, 65)_ = 1.005, *p* = .422), or inactivation x block x sex interaction (*F*_(5, 65)_ = .905, *p* = .483; Figure 9b-c), so sexes were again combined for analyses.

**Figure 9.**
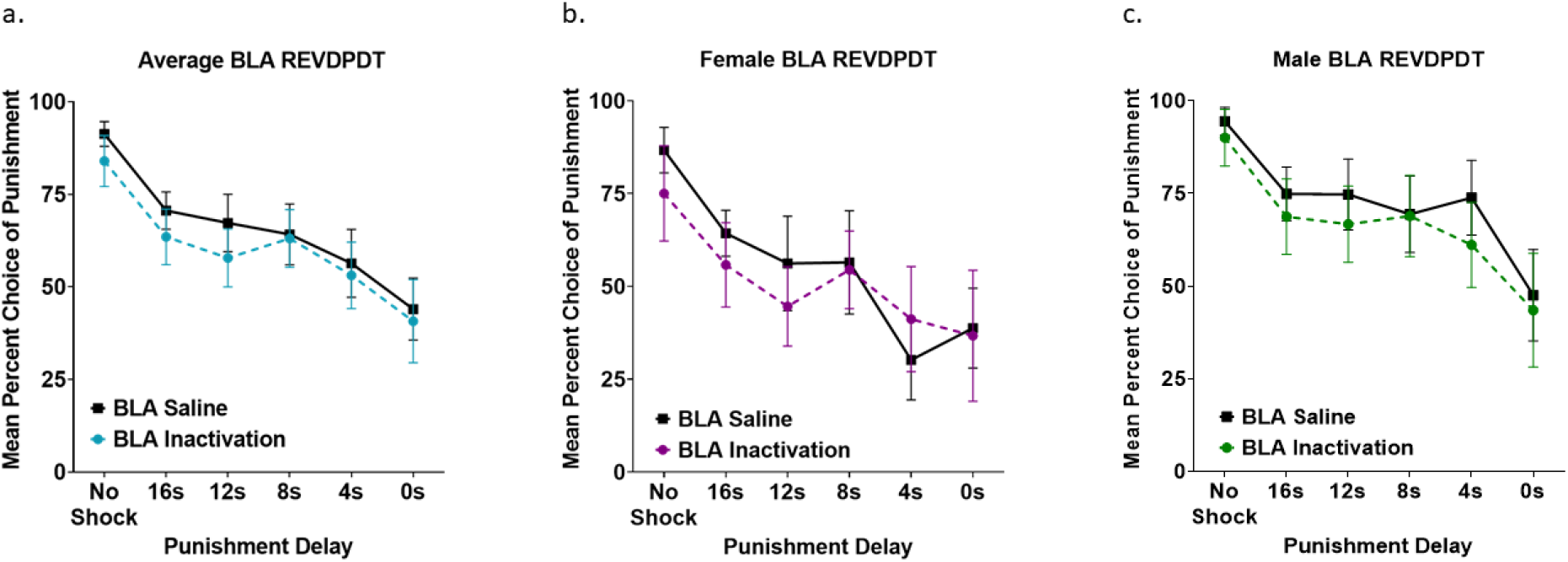
a) BLA inactivation did not affect decision-making during REVDPDT. b-c) Neither females or males were affected by BLA inactivation. All panels display data as mean + SEM.

There was no effect of inactivation (*F*_(1, 13)_ = .444, *p* = .517) or inactivation x block interaction (*F*_(5, 65)_ = .427, *p* = .828; Figure 9a) on choice of the punished reward. There were also no effects of inactivation for any block (*p* > .05; Table 2b), suggesting that unlike standard DPDT, BLA inactivation had no effects on REVDPDT with descending punishment delays.

As with LOFC DPDT and REVDPDT previously, BLA REVDPDT omissions increased throughout the session (*F*_(2.405, 31.263)_ = 3.943, *p* = .023; Figure 10a). However, BLA inactivation had no effect on omissions (*F*_(1.000, 13.000)_ = .070, *p* = .795). There was also no effect of sex (*F*_(1, 13)_ = 1.660, *p* = .220), inactivation x block interaction (*F*_(2.199, 28.589)_ = 1.082, *p* = .357), inactivation x sex interaction (*F*_(1, 13)_ = .695, *p* = .419), or inactivation x block x sex interaction (*F*_(5, 65)_ = .195, *p* = .963; Figure 10a-c).

**Figure 10.**
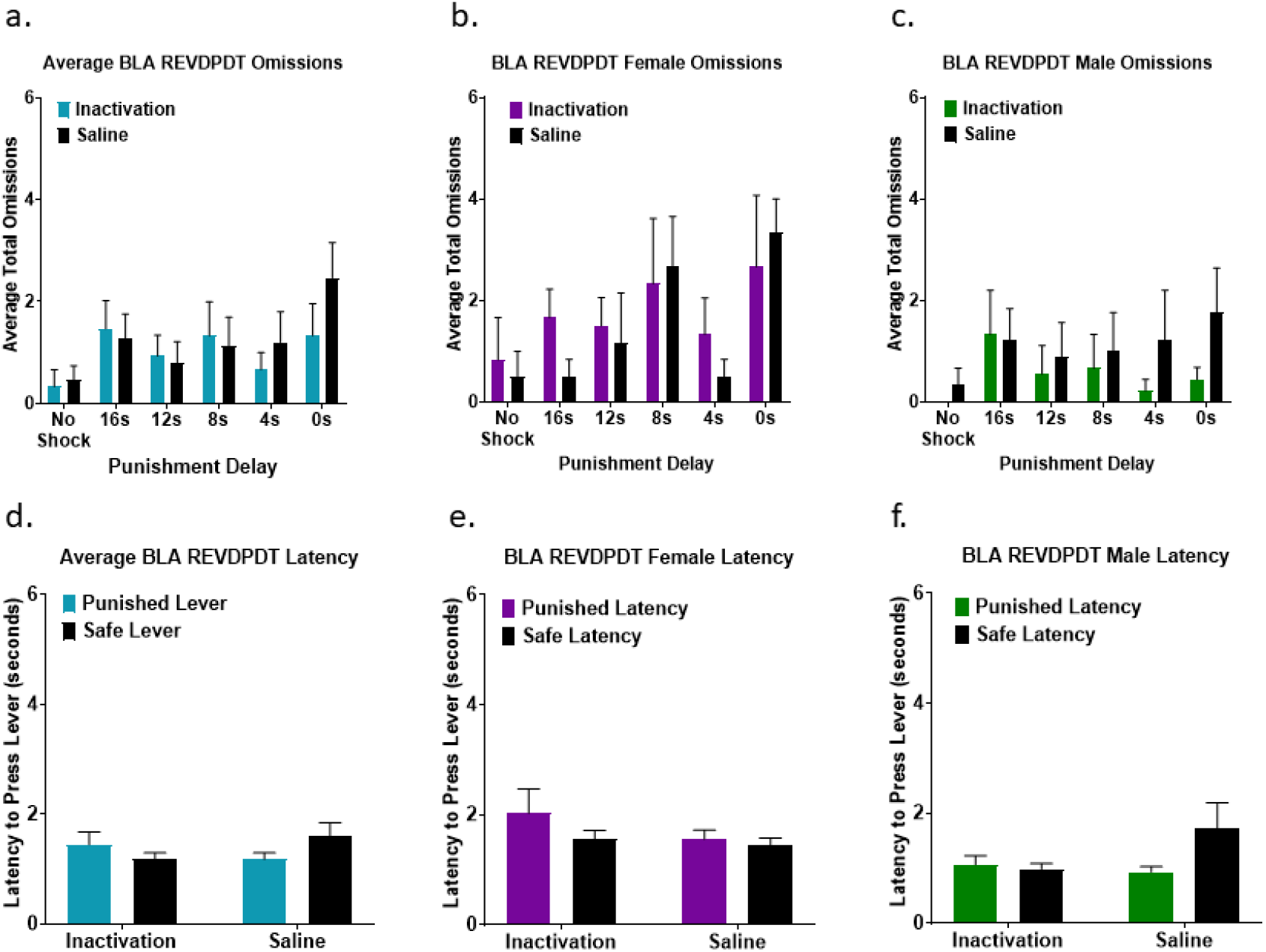
a) BLA inactivation did not affect omissions during REVDPDT. b-c) Females and males displayed a comparable number of omissions. d) Latency to choose between the punished and safe levers was not affected by inactivation. e-f) Females had greater latency to select a lever than males.

Finally, we assessed the effects of BLA inactivation on decision-making latency. There was no main effect of inactivation (*F*_(1, 11)_ = .003, *p* = .960), difference in choice latency between punished vs safe rewards (*F*_(1, 11)_ = .073, *p* = .792), or inactivation x lever type interaction (*F*_(1, 11)_ = 4.435, *p* = .059; Figure 10d). There was a main effect of sex (*F*_(1, 11)_ = 8.871, *p* = .013), such that females took longer to respond than males. There was no sex x lever type interaction (*F*_(1, 11)_ = 2.319, *p* = .156) or sex x inactivation interaction (*F*_(1,11)_ = 2.368, *p* = .152; Figure 10e-f).

## Discussion

While discounting of delayed rewards has been well-studied, little is known about the neural substates underlying delayed punishment discounting. Here we replicated previous findings that rats undervalue punishment preceded by a delay, reflected as increased choice of rewards with delayed compared to immediate punishment. This increased choice of delayed punishment was comparable between ascending and descending punishment delay schedules. LOFC inactivation reduced choice of rewards associated with delayed but not immediate punishment in both male and female rats, suggesting that LOFC contributes to the delay discounting of punishment. BLA inactivation also reduced choice of ascending delayed punishments, but not punishments with descending delays.

### LOFC Regulates Sensitivity to Delayed Punishment

LOFC inactivation reduced choice of rewards with delayed (but not immediate) punishment, indicative of reduced delay discounting. This suggests that LOFC contributes to underestimation of delayed punishment during reward seeking. Further, this is comparable to OFC driving discounting of delayed rewards (Mobini et al., 2002b; Rudebeck et al., 2006), although effects of OFC manipulation vary based on task design and individual differences in impulsivity (Winstanley, 2004; Zeeb et al., 2010). Notably, a population of neurons in OFC signals reduction in value of delayed rewards (Roesch et al., 2006); it is possible that OFC activity signals discounting of impending punishment in similar fashion. However, based on the lack of correlation between delay discounting of reward and punishment (Liley et al., 2019), it is feasible that OFC encodes delayed outcomes differently based on motivational valence.

One explanation for reduced choice of delayed punishment after LOFC inactivation is impaired ability to adapt to changes in delay. This inability to update task contingencies would manifest as a “flattened” discounting curve. However, this is unlikely based on effects of LOFC inactivation during REVDPDT, in which punishment delays decreased throughout the session. As with standard DPDT, LOFC inactivation reduced choice of longer punishment delays (16s) but not immediate or shorter punishment delays, resulting in a “steeper” curve. This verifies that LOFC inactivation does not impair behavioral flexibility in this context.

It is also possible that reduced choice of delayed punishment following LOFC inactivation was not caused by reduced delayed punishment discounting, but by increased overall sensitivity to punishment. However, this is unlikely because LOFC inactivation did not influence choice when punishment was immediate or after a short delay (0-8s). Another possible explanation for reduced large reward choice is that LOFC inactivation impaired magnitude discrimination, as LOFC has been shown to signal reward value (Van Duuren et al., 2008; Simon et al., 2015; Ballesta and Padoa-Schioppa, 2019). This is unlikely because both DPDT and REVDPDT include a punishment-free 1 vs 3 pellet block, during which LOFC inactivation did not influence reward choice.

It is possible that rats performing DPDT are not discounting delayed punishment but are instead unaware of impending punishment due to reduced temporal contiguity between action and outcome, leading to increased choice of options with delayed punishment. OFC has a well-established role in outcome representation (Ursu and Carter, 2005; Mainen and Kepecs, 2009; Panayi et al., 2021); therefore, if choice of delayed punishment was driven by reduced punishment expectancy, inactivation of LOFC would further disrupt expectancy and increase choice of delayed punishment. However, LOFC inactivation here had the opposite effect, reducing choice of rewards with delayed punishment. Therefore, the most plausible explanation for the reduced choice of delayed punishment is reduced punishment delay discounting.

A previous study determined that males discount delayed punishment more than females when shock intensity was comparable for all subjects (Liley et al., 2019). Sex differences in decision-making were not observed here, as shock levels were titrated to avoid ceiling or floor effects for each subject (Orsini and Simon, 2020). Therefore, the lack of sex differences here is a function of task design and does not contradict previous data.

These results suggest that LOFC and BLA regulate DPDT similarly in both males and females. Notably, due to difficulty with task acquisition, the sample size of females was smaller than males. However, due to the lack of sex differences in sensitivity to inactivation, males and females were merged for each experiment. Female rats omitted more trials than males, consistent with previous data showing that estradiol drives avoidance during punishment-based decision-making (Orsini et al., 2021). Inactivation also increased latency for females to make a choice compared to males. Finally, females required almost three times as many sessions as males to acquire DPDT. This is likely attributable to the first exposure to immediate shock driving avoidance of all options (including safe choice) in females. This subsequently increased the time required for females to be exposed to all task parameters, attenuating the overall rate of task acquisition. Alternatively, when the task began with delayed punishment (REVDPDT), females acquired the task as quickly as males. Therefore, beginning training with the option of immediate punishment may cause females to generalize punishment to both levers and completely disengage from the task early in training.

### BLA Inactivation Selectively Reduces Choice of Delayed Punishment

Like OFC, BLA inactivation did not reduce choice of rewards with immediate punishment, but decreased selection of delayed punishment. This is somewhat surprising, as BLA inactivation increased punished choice in both a single choice conflict task and risky decision-making task (Jean-Richard-dit-Bressel and McNally, 2015; Orsini et al., 2015b). It also indicates that BLA inactivation only reduces punishment choice in situations with dynamic punishment delays. The reduced delay discounting of punishment after BLA inactivation is consistent with optical BLA inactivation reducing delay discounting of reward (Hernandez et al., 2019), although BLA lesions increase reward discounting (Winstanley, 2004).

Interestingly, BLA inactivation reduced choice of delayed punishment during ascending punishment delays (DPDT), but not descending delays (REVDPDT). It is possible that decision-making beginning with the option of immediate punishment creates a “high stress” scenario wherein BLA is recruited to influence choice. Conversely, REVDPDT begins with no possibility of punishment, which may reduce BLA involvement in decision-making.

While BLA inactivation reduced choice of the punished reward, we also observed that this avoidance persisted after punishment was removed in the final block of DPDT. It is possible that BLA provides information about reward safety that is no longer available after inactivation, leading to avoidance of the large reward regardless of shock presence. This is supported by evidence that BLA contributes to the processing of safety signals during previously punished scenarios (Ng et al., 2018). The perseverative large reward avoidance could also be explained by BLA inactivation impairing ability to adapt to task changes. However, BLA inactivation during REVDPDT had no impact on choice, suggesting that BLA is not critical for flexibility within a session.

There were no sex differences in decision-making following BLA inactivation, but there were sex differences in other behavioral measures. Omissions were greater in females only during DPDT, suggesting that females are prone to omitting trials when sessions begin with immediate shock. Additionally, there was a trend toward increased omissions as the session progressed after inactivation, suggesting that BLA may contribute to overall motivation or task engagement. Finally, as seen with the LOFC experiments, females had longer latencies than males for both tasks.

The OFC and BLA are densely interconnected (Price, 2007), and both contribute to decision-making informed by reward and punishment (Jean-Richard-Dit-Bressel and McNally, 2016; Orsini et al., 2017). Furthermore, connections between LOFC and BLA are critically involved with encoding/retrieving the incentive value of cues and actions to help guide future decision-making (Groman et al., 2019; Malvaez et al., 2019; Sias et al., 2021). We observed that both LOFC and BLA appear to subserve similar roles in delay discounting of punishment, as inactivation of either region reduced choice of delayed punishment. It is possible that BLA-LOFC work in concert to regulate delayed punishment discounting. However, based on the selective effects of BLA inactivation on DPDT but not REVDPDT, BLA may only be engaged in situations beginning with the threat of immediate punishment. Future experiments will investigate the specific role of this circuit in sensitivity to delayed vs. immediate punishment.

## Conclusions

Insensitivity to delayed punishment is a critical aspect of psychiatric illnesses, during which future consequences are often undervalued in favor of immediate rewards. To our knowledge, this is the first assessment of the neurobiological mechanisms underlying this critical phenotype. These data indicate that LOFC and BLA circuitry may serve as a promising therapeutic target to improve sensitivity to delayed punishment during decision-making.

## Acknowledgements

We thank Mallory E. Udell, Sarah E. Cartwright, Alexandra R. Berwager, Ivan De Jesus Aguilar, Eboni D. Eddins, Lauren M. Clark, and Anna J. Wiener for technical assistance during this project.

